# A Bayesian framework for inferring the influence of sequence context on single base modifications

**DOI:** 10.1101/571646

**Authors:** Guy Ling, Danielle Miller, Rasmus Nielsen, Adi Stern

**Affiliations:** School of Molecular Cell Biology and Biotechnology, Tel-Aviv University, Tel-Aviv, Israel; Department of Integrative Biology, University of California, Berkeley, California 94720, United States of America; Department of Statistics, University of California, Berkeley, California 94720, United States of America; Center for Computational Biology at UC Berkeley (CCB); Edmond J. Safra Center for Bioinformatics at Tel Aviv University

## Abstract

The probability of single base modifications (mutations and DNA/RNA modifications) is expected to be highly influenced by the flanking nucleotides that surround them, known as the sequence context. This phenomenon may be mainly attributed to the enzyme that modifies or mutates the genetic material, since most enzymes tend to have specific sequence contexts that dictate their activity. Thus, identification of context effects may lead to the discovery of additional editing sites or unknown enzymatic factors. Here, we develop a statistical model that allows for the detection and evaluation of the effects of different sequence contexts on mutation rates from deep population sequencing data. This task is computationally challenging, as the complexity of the model increases exponentially as the context size increases. We established our novel Bayesian method based on sparse model selection methods, with the leading assumption that the number of actual sequence contexts that directly influence mutation rates is minuscule compared to the number of possible sequence contexts. We show that our method is highly accurate on simulated data using pentanucleotide contexts, even when accounting for noisy data. We next analyze empirical population sequencing data from polioviruses and detect a significant enrichment in sequence contexts associated with deamination by the cellular deaminases ADAR 1/2. In the current era, where next generation sequencing data is highly abundant, our approach can be used on any population sequencing data to reveal context-dependent base alterations, and may assist in the discovery of novel mutable sites or editing sites.

## Introduction

Single base modifications, which include point mutations and DNA or RNA modifications, are often caused by enzymatic activity. Base alterations can include either standard point mutations or modifications such as cytosine methylation at the DNA level (Cooper and Krawczak 1989), or Adenine to Inosine at the RNA level (Wulff, Sakurai, and Nishikura 2011). For DNA/RNA modifications, specific sequence contexts influence the probability the enzyme will modify a base within this context (Feltus et al. 2006; Lehmann and Bass 2000; Wang et al. 2013). This is reflected by hotspots of mutation in different genomes driven by specific sequence contexts (Aggarwala and Voight 2016; Hodgkinson and Eyre-Walker 2011; Dick G Hwang and Green 2004). One well-known example is that of C->T mutations that occur at high rates in vertebrate genomes by spontaneous deamination of methylated cytosines in CpG positions, i.e. positions in which a Guanine follows a Cytosine (Coulondre et al. 1978; Razin and Riggs 1980). Moreover, specific cellular enzymes belonging to the APOBEC family increase the rate of deamination of Cytosine bases as a means of viral restriction. These enzymes increase the mutation rate of HIV by several orders of magnitude, at specific sequence contexts (Cuevas et al. 2015). Nevertheless, most evolutionary models commonly assume that every position evolves independently. This implies that neighboring positions do not affect the rate of mutation in a given position. Here we present a novel Bayesian method for the analysis of deep population sequencing data, which detects the effect of context on the rate of single base modifications.

Previous efforts for detecting the effects of context on substitution rates were mostly phylogenetic-based methods. In order to keep the number of parameters small and computationally tractable, most such methods consider dinucleotide (Lunter and Hein 2004; Simmonds et al. 2013) or trinucleotide contexts (Blake, Hess, and Nicholson-Tuell 1992; D. G. Hwang and Green 2004). Recently, a novel model for analyzing context dependence in human polymorphisms was suggested (Aggarwala and Voight 2016). However, this model is very high dimensional, and in essence, assumes that a very large number of sequence motifs may influence the rate of substitution. An additional complication is that within the high dimensional space of sequence motifs, there is a high level of correlation between motifs, which could lead to a faulty inference of context effects. For example, the motif CGCXX is contained within the motif XGCXX (where X is any one of the four nucleotides) and this could lead to difficulty in inferring which motif truly has an effect. Here, we develop a model that addresses these problems and includes two main novelties. First, it is tailored for next-generation sequencing (NGS) of populations, data which are becoming more and more abundant. Notably, our method is inspired by ours and other experiments that sequence microbial populations at great depth (Acevedo, Brodsky, and Andino 2014; Stern et al. 2017); typically such experiments result in over 100,000-1,000,000 sequenced viral genomes, with sequencing accuracy that allowed detection of mutations present at a frequency as rare as 10^−6^. The second novelty is that our method explicitly models the assumption that only a few sequence contexts significantly influence the substitution rate, thereby avoiding the combinatorial increase in parameter number with a larger context. This approach relies on the biological motivation that only a limited number of enzymes influence the rate of base alteration and they are often defined by a very specific context. For example, Adenosines in mRNA may be methylated by a methyltransferase enzyme but only in the context RGACU (where R is a purine) (Narayan, Ludwiczak, Goodwin, & Rottman, 1994). There are over one thousand possible contexts in a window size of five (*k* = 5), but since only two of them exert an effect, this is an assumption worthwhile taking into consideration.

When developing the model, we found that the problem of detecting contexts that affect the rate of base alteration bears many resemblances to the problem of quantitative trait locus (QTL) mapping, where the attempt is to detect a limited set of quantitative traits that affect a specific phenotype. As described below, we drew inspiration from QTL models when designing the new model. We tested our method on simulated data that was aimed at mimicking an increase or decrease of mutation rate, in a population of replicating viruses. Our model accurately captured these changes in the simulated data. We next analyzed data from next-generation sequencing experiments of polioviruses and describe how our method captures intriguing biological signals.

## Methods

### Context-dependent base alteration model

For the sake of simplicity, we hereby refer to a base alteration as a mutation. This is often convenient since a base alteration may be captured in next-generation sequencing experiments as an observed mutation (for example, Adenine to Inosine alterations are observed as Adenine to Guanine mutations following sequencing (Levanon et al. 2004)). Let the full sequence (e.g., the genome) we are interested in being denoted as *G* = (*g*_1_,…,*g_n_*), where *g_i_*, ∈ {*A,C,G,T*} and |*G*| = *n*. We denote the *k*-long context of a focal position, *i*, as the 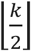 nucleotides flanking the position, i.e., the sequence 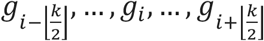, where *k* is assumed to be an odd number. For example, for *k* = 3 and *G* = *AGGAT* there are three distinct contexts, *AGG, GGA*, and *GAT* (Figure 1).

**Figure 1.**
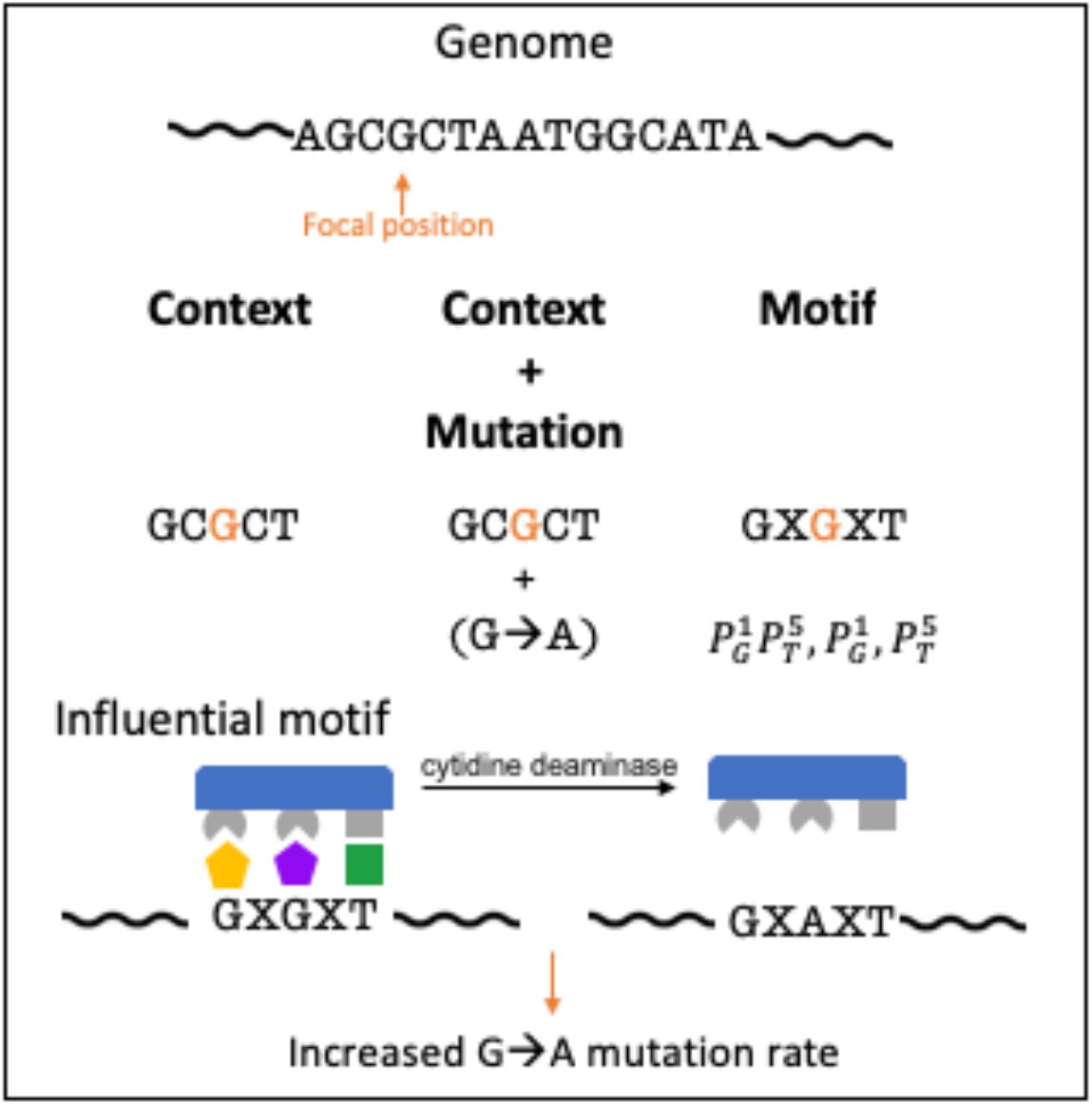
Model definitions. For the labeled focal position G, we present its genomic context under *k*=5, and illustrate the context, the context together with a mutation, and a possible motif embedded in this context. Motifs are associated with indicator functions labelled as 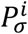 that will be set to 1 if and only if the *i*th position of the context is *σ* (see main text). We exemplify how enzymatic activity operating on a specific sequence context may result in an increase in the mutation rate at this context.

We can further decompose sequence contexts into motifs (which will be the features in our feature selection algorithm). In the example above, a possible motif would be *AGX* (where *X* is any of the four nucleotides). Formally let us define a motif as a context pattern and denote 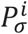 as a binary indicator function for the presence of character *σ* at the *i*th position of the context. In our example, *AGG* is a context, and the indicator for the motif *AGX* 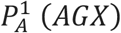 will be set to 1 because the nucleotide in position 1 is indeed *A*. Similarly, the indicator for 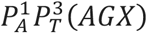 will be set to 0 because the 3^rd^ position in the motif *AGX* is *G* and not *T*. For *k*=1, there are 4 motifs (the 4 possible nucleotides), for *k*=3 there are 3 × 4 motifs consisting of one nucleotide, 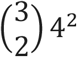 motifs consisting of two nucleotides, and 4^3^ motifs consisting of 3 nucleotides resulting in a total of 120 motifs. In general, for a *k*-long context, there are 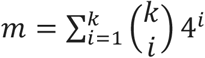 motifs (Figure 1 and Table 1). Next, we consider a mutation operating on a context, *μ* = (*a, b*), to be an ordered 2-tuple, where *a,b* ∈ {*A,C,G,T*},*and a*≠*b* (Figure 1).

**Table 1.**
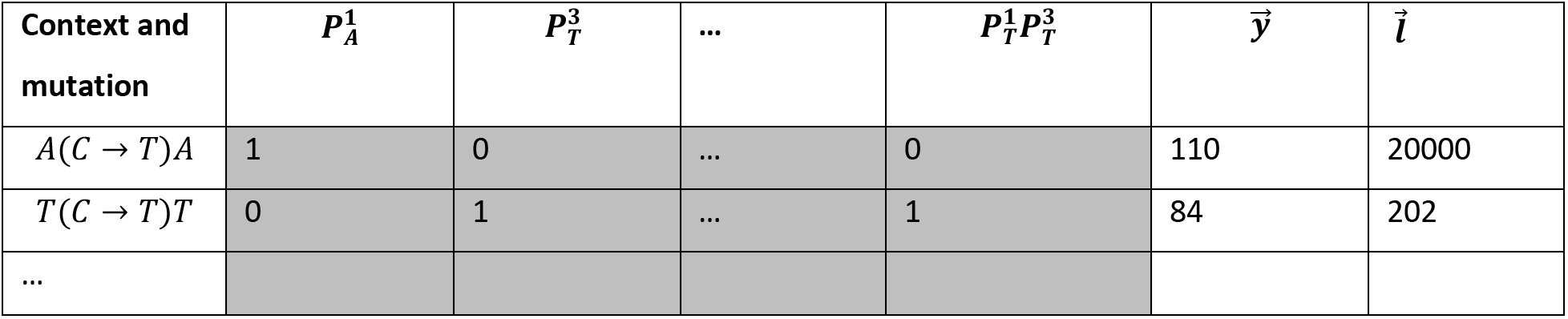
An example of sequencing data and associated breakdown into context, motifs, and counts. The main variables *X* are shaded in grey, representing the indicator functions *P* that dictate presence/absence of motifs within a context. In the first row of this example, we observe C➔T mutations within the context *ACA* in 110 of the individuals in the population sequenced, out of a total of 20,000 sequences (reads) covering the two focal positions.

| Context and mutation | $P_A^1$ | $P_T^3$ | ... | $P_T^1 P_T^3$ | $\vec{y}$ | $\vec{l}$ |
| --- | --- | --- | --- | --- | --- | --- |
| $A(C \rightarrow T)A$ | 1 | 0 | ... | 0 | 110 | 20000 |
| $T(C \rightarrow T)T$ | 0 | 1 | ... | 1 | 84 | 202 |
| ... |  |  |  |  |  |  |

Now, let 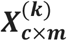 be a design matrix that indicates, for each context of length *k* that appears in the genome and a mutation (*a, b*), the motifs it contains such that:

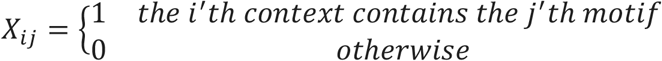

where *c* is the number of all unique contexts of genome *G* and their corresponding observed mutations, and *m* is defined above as the number of possible motifs (Table 1). Notably, a given context may be present more than once in a genome, or not be present at all.

Given a population of similar genomes that has been sequenced we can define the vector 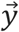 to be the empirical count for each mutation at a given context, so that *y_i_* is the observed mutation count for the *i*th context with mutation *μ* = (*a, b*), i.e, the number of mutations observed at all focal positions sharing the same context. We define 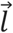 as a vector of the total observed number of sequences at each context so that 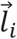 is the number of observed sequences covering all focal positions at the *i*th context (i.e., the sequencing coverage at position *i*). Table 1 illustrates an example showing *X*, 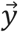, and 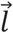 for a context of size *k*=3 for simplicity.

### Model

We will use a logistic regression model with the latent variables *α* ∈ *R,σ* ∈ *R*^+^, 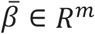, and 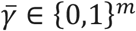 (where *m* is the number of all possible motifs), which relates motifs in a context to the probability of observing a mutation in the context, defined as 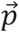. We assume that *y_i_*~*Binomial* (*l_i_,p_i_*) and 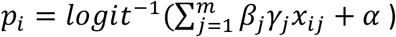. The introduction of the 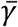 indicator vector allows us to better penalize models with a large number of influencing motifs, i.e. it will allow us to use shrinkage in the model to avoid overfitting. Thus, our logistic regression model is defined by 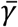, a vector of indicator variables indicating whether the *ith* variable contributes significantly to the model or not, 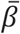, the vector of regression coefficients that increase or decrease the probability of a mutation in a context, and *α*, the baseline mutation rate in the absence of any context effect. We use the properties of the 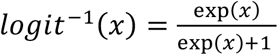 which maps *logit: R* → (0,1) to guarantee that *p_i_*, is always a valid probability.

### Inference

We will use Bayesian variable selection to estimate 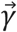. Our approach is similar to classical methods used for quantitative trait locus (QTL) mapping (Yi 2004). These methods aim to identify correlations between a set of genetic markers (e.g., single nucleotide polymorphisms, SNPs) and a continuous phenotype (Yi 2004). There is usually a very large number of SNPs (which take the role of features) in a sample and a much smaller number of samples. Moreover, often the different features are correlated, mainly due to linkage in a genetic cross or linkage disequilibrium in outbred populations. In our case, the motifs are the features and the number of possible motifs is often much larger than the number of mutations observed in a given genome, especially when considering microbial genomes that tend to be relatively small. Furthermore, there is also a strong correlation between the different motifs (features), since these may be nested within each other or may be mutually exclusive. We address these problems by assigning a prior distribution to each latent variable. The model is too complex to allow analytical solutions, but we can infer the posterior distribution using a Markov Chain Monte Carlo (MCMC) algorithm.

### Posterior probability calculations

The posterior probability of the four latent variables can be written as:

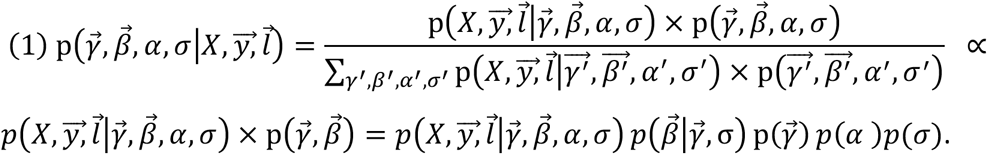

The expressions in the numerator can easily be computed when values for the latent variables are given: 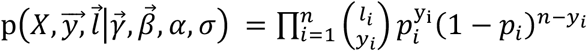, where *p_i_* is defined above. On the other hand, calculating the sum in the denominator is intractable for all possible combinations of 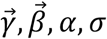. We thus chose to use MCMC to traverse the posterior distribution using a Metropolis-Hastings method (Statistician and Nov 2007).

### Prior probability specification

#### Prior for 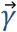

We define a simple Bernoulli prior probability distribution for each element of 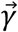 so that 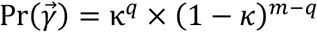 for some hyper-parameter *k*, where *q* is the number of ‘1’ entries in 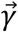. Taking *κ*= 0.5 defines a prior that gives equal weight to any 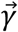. In practical estimation, the collinearity between the predictors can lead to instability. The collinearity for this problem has been described previously as a “dilution” effect (George 2010). For example, if 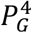 is a feature that increases the mutation rate for *C* → *T* mutations, naïve estimation of effects by counting might also find that 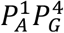 significantly increases the mutation rates, as the 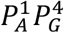 motif contains the 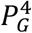 motif. To address this problem, we add a penalty to the prior for 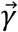. We use a method based on Determinantal Point Process (DPP) that has been shown to be effective in other Bayesian variable selection problems (Kojima and Komaki 2014; Ročková and George n.d.). According to the DPP method, the prior is weighted by powers of the determinant of the correlation matrix, 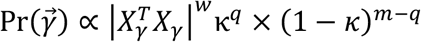, where *X_γ_* is a *n× q* design matrix including only the columns 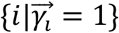 from the original matrix *X* and *w* ∈ *R*^+^ is a weight factor. The weighting provides a computationally tractable approach for mitigating the effect of the dependencies between the features. If all features are completely independent 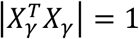, whereas in the case of full dependency (collinearity) between at least one pair of vectors the matrix will be singular and 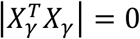.

#### Prior for β

We define the prior probability for the regression coefficients, 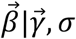, as: 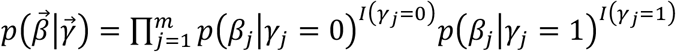, where (*I*(·) is the indicator function, *p*(*β_j_|γ_j_* = 0) = *N*(0,*σ*^2^), and *p*(*β_j_|γ_j_* = 1) = *N*(0,*C*^2^*σ*^2^) for some variance *σ*^2^ and some constant *C*. It is possible to either infer *σ*^2^ from the data or to define it as a constant; here we chose to infer it from the data. Notably when *γ_j_* = 0, the regression coefficient is undefined and *β_j_* is unmeaningful.

#### Prior for α

*α* represents the mean rate of base modification, which we refer to here as the base mutation rate. We assume α is normally distributed 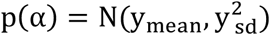 and use the empirical mean and variance inferred from the data, thus 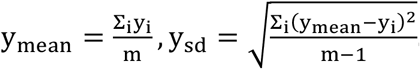.

#### Prior for σ^2^

*σ*^2^ can be set as a hyperparameter or can be inferred from the data with a variety of possible priors. A simple prior for *σ*^2^ is the uniform prior *σ*^2^~ *U*(*low, high*), where *low* and *high* are set arbitrarily.

#### MCMC implementation

Our MCMC algorithm works as follows:

1. We start from an arbitrary point (*β*_0_, *γ*_0_, *α*_0_, *σ*_0_).
2. We then define a transition kernel, i.e., the set of probabilities for proposing a new parameter value given the previous value. Let the parameters in the *i*’th step be *β_i_, γ_i_, α_i_, σ_i_*. A new set of parameters 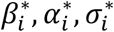 are sampled based on Gaussian distributions for each variable: 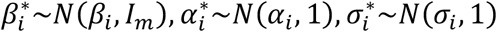. For *γ_i_*\* (here *i* denotes the algorithm step and not the vector entry) we can take one of two possible approaches. The first is simply randomly selecting entries in *γ_i_* and flipping them (from 1 to 0 and vice versa) and thus creating a 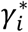 with Hamming distance 1 from *γ_i_*. The second approach which allows more diversity in the 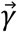 steps is to assume 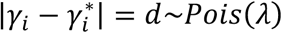 for some *λ* and randomly choose *d* entries and flip them. In this analysis, we used the second approach since it mitigates the problem of local optima.
3. Then we accept 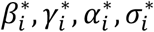 with probability 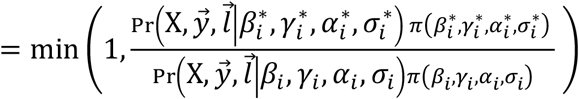, where *π*(·) is the prior.
4. After the chain has completed a pre-determined number of iterations (burn-in), we use the ergodic averages of each parameter to approximate the posteriors.

### Moran model for simulating context dependence

To simulate viral sequences where the rate of mutation depends on the context, we used an adaptation of the 4-allelic Moran model for each position of the genome previously described for simulating the evolution of cancer cells (Zhu et al. 2011). This model is a continuous time birth-death model. Usefully, this model takes into account large population sizes and allows for different mutation rates, both of which are relevant in our case. We simulate each position separately, while taking into account the context of the position as described below. Each allele is one of the four nucleotides.

Let *N* be the population size. For a given position *i*, at time *t* we can define a vector 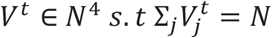. Thus, each entry 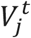 is the number of genomes with the *j^th^* allele in the *i’th* position. We initialize

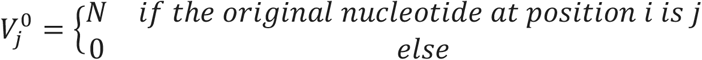

meaning that we start the simulations from a homogenous population of genomes defined by the original sequence. When an individual of type *j* dies it is replaced by randomly selecting a parent from the *N* options and “cloning” it. Notably, the model we used (Zhu et al. 2011) also defines for each phenotype *j* a different fitness value (1 + *s_j_*) and different mutation rates *μ_jj_’* for each *j* ≠ *j*’ that changes the allele of the individual. Here, we assume neutrality for all mutations, and hence set *s_j_* = 0 for every *j*. In order to model the influence of context, we assume there are different *μ_jj_’* for different sequence contexts. For each position at a given context, 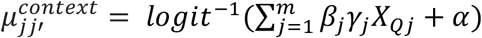, similar to the way context was assumed to affect mutation rates in our model above. The evolutionary process defined above is a continuous time Markov chain. At time *t* the state of the chain is 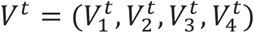. For a 4-allelic model, we have 12 possible transitions that can change the state of the chain (*V_j_* → *V_j_*, for all *j* ≠ *j*’). For a transition of type *k*, = 1,…,12, involving a change from *j* → *j*’ the relative transition rate is

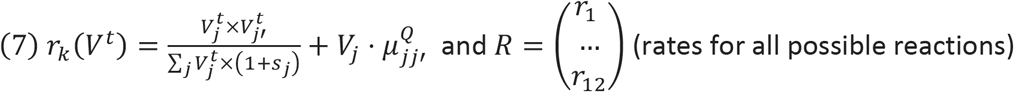

After defining the rate of the *k’th* substitution rate, let’s assume it’s a reaction that changes the vector *V^t^* by adding one to 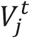 and subtracting one from 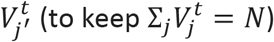. We can say that the substitution changes the vector *V^t^* by adding *ξ^k^* to it, where:

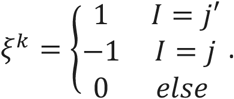

We define a matrix *S*_4×12_ = 〈*ξ*^1^|…|*ξ*^12^〉 and vector 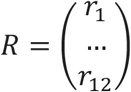 and we can show that 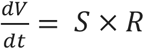

After defining the Markov chain, we can simulate it by using the stochastic simulation algorithm of Gillespie (Gillespie 1977), which takes as parameters the initial conditions, the matrix *S* and the rate vector *R* and outputs a possible chain *V^t^* for a given time *t*.

### Simulating noisy frequencies

In order to simulate mutation frequencies that underwent both sampling and sequencing errors, we performed two steps: we began with the set of simulated frequencies described above which we assumed to be our true frequencies and assumed *N*=1,000,000 genomes. In the first step, we applied binomial sampling on the frequencies and sampled 100,000 genomes for sequencing. Second, we applied a transition probability matrix *A* that introduces sequencing errors, where *A_ij_* is the probability of a sequencing error from one base *i* to base *j*. Error probabilities were defined based on our characterization of errors present in Illumina-based sequencing (Gelbart et al. 2018).

## Results

### Simulated datasets

In order to verify the performance of our method, we simulated population next-generation sequencing data. Our aim was to mimic evolving populations of viruses where rare mutations are often observed widely across the genome. Accordingly, our simulations mimicked population sequencing of oral poliovirus (Stern et al. 2017) with a genome length of ~7,500 bases at a sequencing depth of ~100,000 reads per locus. Using an adaptation of the Moran model (Methods), we introduced an increased or reduced mutation frequency based on *k*=5 contexts. Our simulations began with a homogenous population of genomes. Fourteen generations of mutations only (no selection) were simulated, with a population size *N*=10^5^ and mutation rate *u*= 10^−5^. In each of 500 simulated datasets, we introduced a different number (between zero and three) of motifs influencing the mutation rate of a specific mutation type. We then used our inference framework to infer which were the influential motifs in each of the simulations. We define a motif as one that influences the mutation rate if the expectation of the empirical gamma posterior distribution exceeded some pre-defined threshold *t*, i.e. we will reject the null hypothesis (*γ_i_* = 0) if 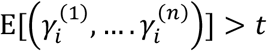 for a given gamma posterior 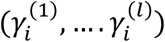.

Table 2 summarizes the results of the simulations given different thresholds of *t*. All in all, our accuracy rate was very high across all thresholds tested, demonstrating the power of our approach to accurately detect the influence of a motif on the rate of mutation. Our low false positive rate, together with a high true negative rate across all thresholds demonstrates that the method has high specificity, i.e., we correctly reject non-influential motifs. However, we are not able to detect all the influential motifs, as demonstrated by the somewhat low true positive rate.

**Table 2.** Inference results on simulated data: TPR (True Positive Rate), TNR (True Negative Rate) and ACC (Accuracy) for each of the different thresholds (*t*).

| $t$ | $TPR$ | $TNR$ | $ACC$ |
| --- | --- | --- | --- |
| 0.4 | 0.83 | 0.96 | 0.95 |
| 0.5 | 0.83 | 0.96 | 0.95 |
| 0.6 | 0.83 | 0.964 | 0.96 |
| 0.7 | 0.83 | 0.964 | 0.96 |
| 0.8 | 0.83 | 0.964 | 0.96 |
| 0.9 | 0.83 | 0.964 | 0.96 |
| 1 | 0.82 | 0.965 | 0.961 |

We illustrate our analysis with one of the simulated datasets. The motifs originally simulated as altering the mutation rate were 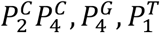, and their corresponding *β* coefficients were −2,2,1, respectively. We then applied the method for 3 × 10^6^ MCMC steps and plotted the posterior distributions and the MCMC trace of the *β* coefficients in Fig. 1. Our posterior distributions show we manage to accurately capture the simulated *β* coefficients.

### Introducing noise into the analysis

In our simulations above all sites were simulated as neutral, and the observed frequencies were assumed to be the true frequencies. However, in real biological data, these two assumptions will likely be problematic. For one, there is no set of genomic sites known to be completely neutral. Typically, synonymous sites are assumed to be neutral, yet a subset of these sites may be under selection (Chamary, Parmley, and Hurst 2006). Moreover, observed mutation frequencies will be affected by sampling and by sequencing errors and thus will likely deviate from true mutation frequencies. Thus, in order to test how our method fares with noisier data, we applied two different “noising” schemes.

We first tested the method when a proportion of sites deviate from neutrality. We simulated 500 datasets with a varying number of loci under selection. Selection coefficients were sampled based on a distribution of fitness effects from an empirical dataset of RNA viruses (data not shown). The accuracy of inference was examined as a factor of *β*, the proportion of non-neutral sites, and sequencing noise. Setting *β* to zero yielded no context effect, as 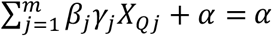, with the area under the curve (AUC) equal to 0.5 as expected. As |*β*| increased, the higher the probability of a mutation in a given context, and accordingly the power of the model rose, and motifs were efficiently identified (Fig. 2A). In fact, for a very large *β* the effects of noise (both sampling and the % of non-neutral sites) were very much mitigated (Fig. 2B-C).

**Figure 2.**
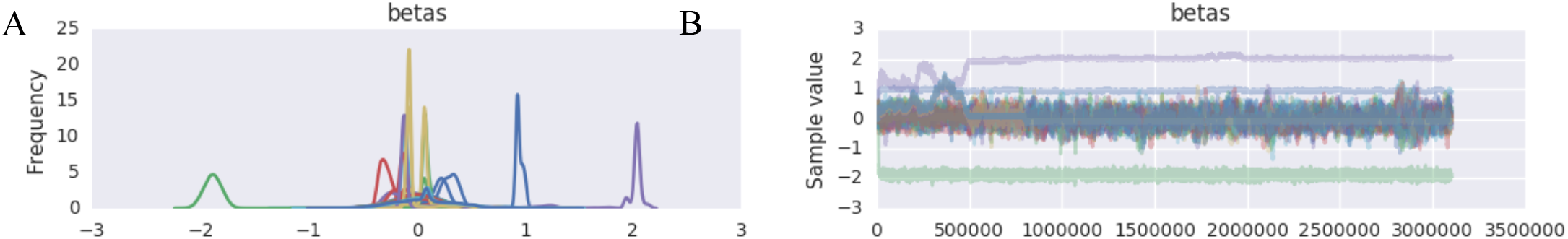
MCMC posterior distribution results for one simulated dataset: (A) Posterior density of *β* values show peaks over the three true values, (B) trace of *β* values show convergence to true values.

**Figure 3.**
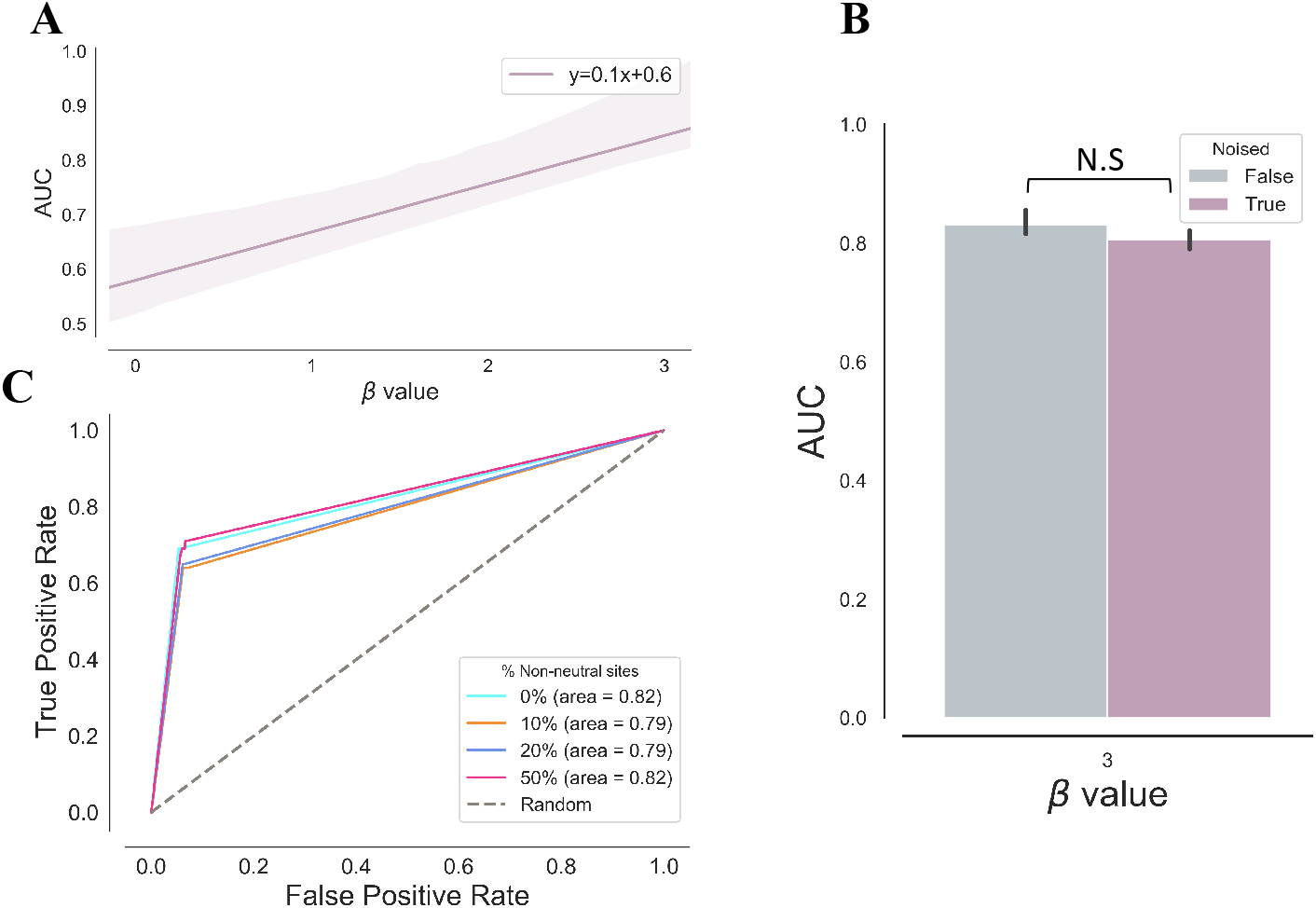
Analysis of 500 noisy context simulations for different values of *β*, across different proportions of non-neutral sites. (A) The change in AUC as a factor of increasing *β* with a 95% confidence interval shown in the violet shading. (B) AUC comparison of simulations with and without noise as defined in the main text. (C) Receiver operating characteristic (ROC) for noisy simulations with *β* = 3.

### Oral polio vaccine virus experimental evolution dataset

To further examine the biological application of our method we applied it to oral poliovirus 2 (OPV2) sequencing data (Stern et al. 2017). The experiment was designed to record the mutation frequencies of OPV as it was serially passaged in tissue culture. Notably, the very high sequencing depth of this experiment, spanning 10^5^ – 10^6^ reads per position, combined with very high mutation rates of the virus (spanning 10^−4^ – 10^−5^ mutations per base per replication cycle), made these data perfect for our model. In order to rule out the effects of selection that can easily mimic the effects of increased or decreased mutation rate, our analysis focused only on synonymous mutations that are mostly (although not always) neutral. We further focused only on transition mutations, since transversions are less frequent and hence inferred with less reliability. We analyzed both the last and seventh passage, which corresponded to fourteen viral replication cycles.

Table 3 presents the resulting motifs detected in passage 7, given a threshold *t* = 0.7 for the expectation of the gamma posterior distribution. Intriguingly, many of the motifs detected are compatible with editing by the enzymes hADAR1 or hADAR2 (Eggington, Greene, and Bass 2011). Both enzymes edit Adenine to Inosine, which is detected as an A➔G mutation, and prefer A or U (T) upstream to the edited A. Since polioviruses copy both the positive and negative RNA strand synthesis in the cell (Schulte et al. 2015), a T➔C mutation on the reverse-complement negative strand will be read as an A➔G on the positive strand (which is the reference genome against which all reads are mapped). Accordingly, table 3 shows enrichment for A➔G and T➔C mutations as compared to the composition of the OPV2 genome (p < 0.001, Fisher exact test).

**Table 3.**
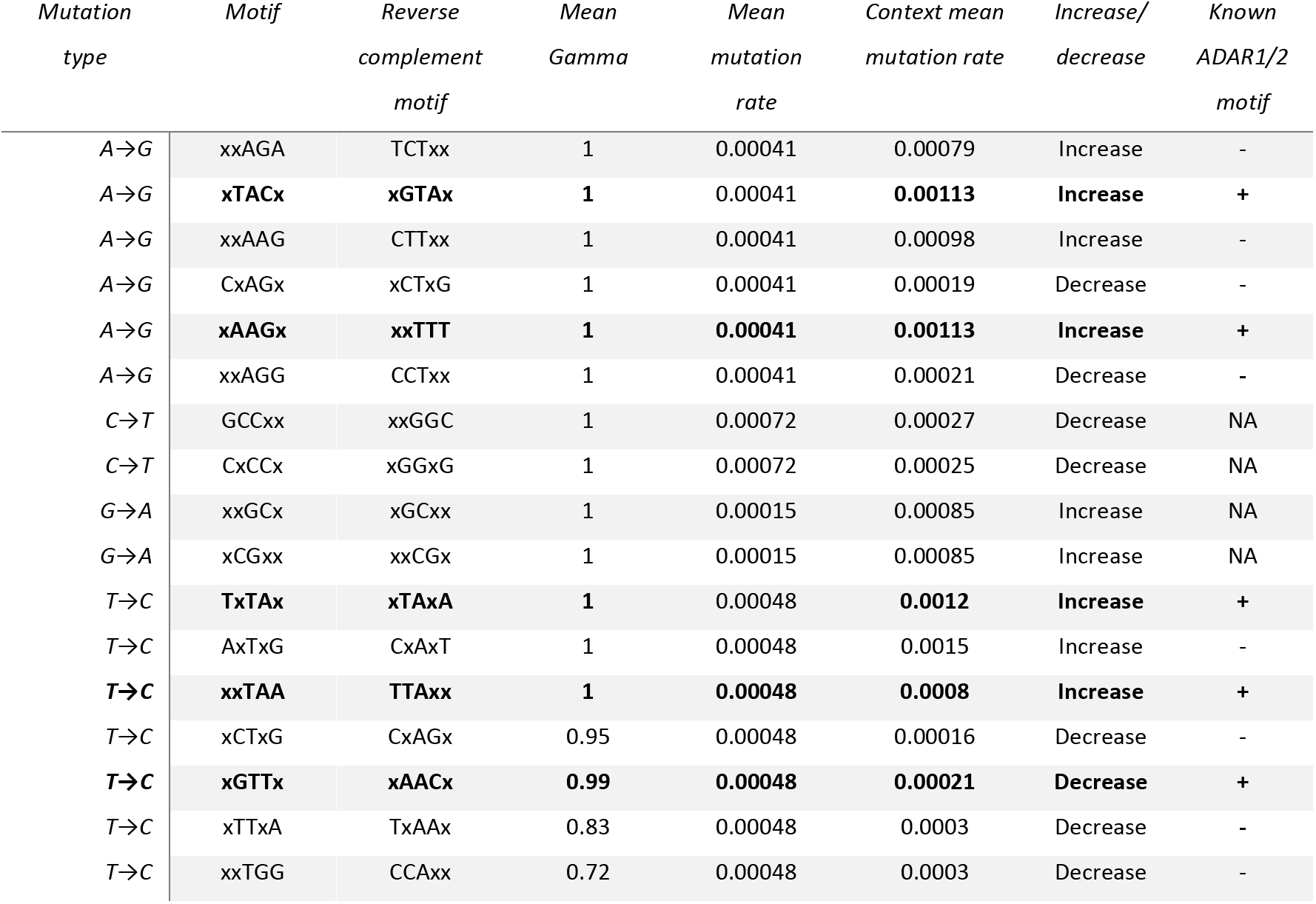
Motifs affecting mutation rates detected in empirical data of passage 7 of OPV2.

## Discussion

We developed here a novel approach for the detection and evaluation of sequence context on mutation rates. Our prime motivation was to develop a method that is able to analyze high-resolution deep sequencing datasets, where the depth of sequencing scale with the mutation rate. One of the main challenges in these datasets is the high dimensionality of the data when accounting for sequence context. Thus, one of the main novelties of the approach herein is that it takes into account the fact that the number of sequence motifs that affect mutation rates is likely much smaller than the number of possible motifs. We conclude that the method provides highly accurate results on simulated data. In particular, we precisely identify the motifs which are not influenceable, regardless of the prior distribution we use (DPP or binomial) and the threshold we impose. Our remarkably low false positive rate promotes high confidence in inferring influencing motifs; however, we also fail to detect some true motifs, suggesting that our method is conservative, a trait which is desirable in such a method.

Having said that, we would like to recognize some limitations that could restrain the effectiveness of the inference using our method. The assumption of mutation neutrality is essential to the analysis as point mutations which strongly deviate from the neutrality might promote the appearance of a non-existing context effect. That caveat is diminished for highly influential motifs (large beta), however, in empirical data the effect of a motif may not be so strong, as we show for the poliovirus data. The more data that is available (e.g., the more synonymous sites analyzed), it is likely that the effect of a handful of non-neutral sites will be diminished.

Our tests have also shown that the method is sensitive to the selection of hyperparameters and specifically to the selection of the *κ* hyperparameter which controls the penalty for the introduction of an influential motif to the model. Using different values may yield slightly different results (data now shown), and it is best to calibrate this parameter by first analyzing simulated data that is similar to the empirical data at hand.

To summarize, we have developed a robust method that has the potential and the strength to identify influential sequence contexts, and this may shed light upon the underlying mechanisms of both polymerases and other enzymes that render genetic modifications. The method is flexible, compatible with a wide variety of applications and data sets, and should be fairly easily executed for the analysis of NGS data. While the method was designed with our recent experiments of virus populations in mind, it is also generic enough for any other type of NGS data.

The python code for the method is publicly available via GitHub under https://github.com/SternLabTAU/context

## Acknowledgments

We would like to thank Karin Mittelman and Shiran Abadi for their constructive suggestions during the writing of this paper. This study was supported in part by a fellowship to GL and DM from the Edmond J. Safra Center for Bioinformatics at Tel-Aviv University, by funding from the Israeli Science Foundation (1333/16) to AS, by funding from Raymond and Beverly Sackler Fund to AS and RN, and by funding by the Koret-UC Berkeley-Tel Aviv University Initiative in Computational Biology and Bioinformatics to AS and RN.

